# A cnidarian phylogenomic tree fitted with hundreds of 18S leaves

**DOI:** 10.1101/2022.10.03.510641

**Authors:** Melissa B. DeBiasse, Ariane Buckenmeyer, Jason Macrander, Leslie S. Babonis, Bastian Bentlage, Paulyn Cartwright, Carlos Prada, Adam M. Reitzel, Sergio N. Stampar, Allen G. Collins, Marymegan Daly, Joseph F. Ryan

## Abstract

Cnidarians are critical members of aquatic communities and have been an experimental system for a diversity of research areas ranging from development to biomechanics to global change biology. Yet we still lack a well-resolved, taxonomically balanced, cnidarian tree of life to place this research in appropriate phylogenetic context. To move towards this goal, we combined data from 26 new anthozoan transcriptomes with 86 previously published cnidarian and outgroup datasets to generate two 748-locus alignments containing 123,051 (trimmed) and 449,935 (untrimmed) amino acids. We estimated maximum likelihood phylogenies for both matrices under partitioned and unpartitioned site-homogeneous and site-heterogenous models of substitution. We used the resulting topology to constrain a phylogenetic analysis of 1,814 small subunit ribosomal (18S) gene sequences from GenBank. Our results confirm the position of Ceriantharia (tube-dwelling anemones), a historically recalcitrant group, as sister to the rest of Hexacorallia across all phylogenies regardless of data matrix or model choice. We also find unanimous support for the sister relationship of Endocnidozoa and Medusozoa and propose the name Operculozoa for the clade uniting these taxa. Our 18S hybrid phylogeny provides insight into relationships of 15% of extant taxa. Together these data are an invaluable resource for comparative cnidarian research and provide perspective to guide future refinement of cnidarian systematics.

## Introduction

Cnidarians have been evolving independently from other animals for at least 600 million years (Erwin 2015; Dohrmann and Wörheide 2017; McFadden, et al. 2021) and have diversified into an astonishingly wide assemblage of forms, including hard and soft corals, anemones, siphonophores, hydroids, jellyfish, and myxozoan parasites. Cnidaria consists of 12,153 extant, accepted species (WoRMS Editorial Board 2022) and approximately 8,000 additional predicted species (estimated from Appeltans, et al. 2012). These diverse species form a well-supported clade and are united by their ability to produce stinging cells called nematocytes (Collins, et al. 2020). The species richness of Cnidaria, the ecological importance of many of its species, and its phylogenetic position as sister to Bilateria, has made Cnidaria the focus of a range of basic biological research questions. As such, the long-standing goal of establishing a complete cnidarian tree of life is becoming more urgent, but also more tractable, as maturing sequencing technologies allow for the collection of more phylogenetic characters from more species.

There is a rich history of research involving cnidarians, with centuries of studies on topics such as regeneration (e.g., Trembley 1744; Zoja 1895; Zeleny 1907), embryogenesis (e.g., Murbach 1896; Hargitt 1904), coral reef formation (e.g., Darwin 1851), life history (e.g., Sars 1829), physiology (e.g., Romanes 1880), systematics (e.g., Müller 1862), and morphology (e.g., Hargitt 1901). Scientific interest in these animals has not waned over time. Research on broad biological questions using cnidarians as a focal system continues in all fields of biology, with striking recent examples including studies of allorecognition (Karadge, et al. 2015), biogeography (Martínez, et al. 2010), biomechanics (Hamlet and Miller 2014), circadian clock (Peres, et al. 2014), development (Helm, et al. 2013), early animal evolution (Martin, et al. 1997; Collins and Valentine 2001; Gröger and Schmid 2001; Bebenek, et al. 2004), evolutionary novelty (Babonis, et al. 2016), genomics (Putnam, et al. 2007; Chapman, et al. 2010; Leclère, et al. 2019), germ cell evolution (Extavour, et al. 2005; Chen, et al. 2020), global change (Hoegh-Guldberg 1999; Bellwood, et al. 2004), human health (e.g., Miller, et al. 2005; Sullivan and Finnerty 2007), life history (Sanders and Cartwright 2015), natural products (Jouiaei, et al. 2015; Mariottini and Grice 2016), neurobiology (Grimmelikhuijzen, et al. 2004; Marlow, et al. 2009), regeneration (Chera, et al. 2009; Bradshaw, et al. 2015), stem cell biology (Gahan, et al. 2016; Siebert, et al. 2019), symbiosis (Davy, et al. 2012; Lehnert, et al. 2012; Newkirk, et al. 2018; Gault, et al. 2021), venom (Macrander, et al. 2015; Macrander, et al. 2016; Klompen, et al. 2020), and vision (Picciani, et al. 2018). This growing community of researchers and an expanding taxonomic breadth applied to a diversity of questions (e.g., He, et al. 2019), underscores the importance of an accurate and comprehensive cnidarian tree of life.

Early efforts to reconstruct the phylogeny of Cnidaria emphasized broad scale patterns, including work by Siddall et al. (1995), who used 18S sequence data to demonstrate that Myxozoa belonged to Cnidaria after Smothers et al. (1994) showed them to be metazoans. Bridge et al. (1995) combined 18S and 16S sequence data with morphological characters to test class-level relationships within Cnidaria. Many multi-locus studies followed, including those that used some combination of nuclear ribosomal (e.g., 5S, 18S, 28S, ITS) and mitochondrial genes (e.g., 12S, 16S, COI, COIII) to resolve relationships within individual cnidarian lineages [**Actiniaria:** (Geller and Walton 2001; Daly, et al. 2008; Gusmão and Daly 2010; Rodríguez, et al. 2014; Grajales and Rodríguez 2016; Larson and Daly 2016; Daly, et al. 2017; Gusmão, et al. 2019; Titus, et al. 2019; Gusmão, et al. 2020; Gusmão and Rodríguez 2021); **Antipatharia:** (Brugler and France 2007; MacIsaac, et al. 2013; Bo, et al. 2018; Barrett, et al. 2020; Horowitz, et al. 2020); **Anthozoa:** (Berntson, et al. 1999); **Ceriantharia:** (Stampar, et al. 2012; Stampar, et al. 2014); **Cubozoa:** (Won, et al. 2001; Bentlage, et al. 2010); **Hydrozoa:** (Cartwright, et al. 2008; Collins, et al. 2008; Nawrocki, et al. 2010; Nawrocki, et al. 2013; Maronna, et al. 2016; Cunha, et al. 2017; Bentlage, et al. 2018; Mendoza-Becerril, et al. 2018; Bentlage and Collins 2021); **Myxozoa:** (Jiménez-Guri, et al. 2007; Evans, et al. 2010; Bartošová and Fiala 2011; Li, et al. 2020); **Octocorallia:** (McFadden and Van Ofwegen 2013; McFadden, et al. 2017; García-Cárdenas, et al. 2020; Sánchez, et al. 2021; Untiedt, et al. 2021; Watling, et al. 2022); **Scleractinia:** (Romano and Cairns 2000; van Oppen, et al. 2001; Fukami, et al. 2008; Huang, et al. 2009; Barbeitos, et al. 2010; Huang, et al. 2011; Stolarski, et al. 2011; Arrigoni, et al. 2012; Prada, et al. 2014; Luzon, et al. 2017; Cowman, et al. 2020); **Scyphozoa:** (Bayha, et al. 2010; Bayha, et al. 2017); **Staurozoa:** (Collins and Daly 2005; Miranda, et al. 2010; Miranda, Hirano, et al. 2016); **Zoantharia:** (Swain 2010; Sinniger, et al. 2013; Montenegro, et al. 2016; Kise, et al. 2019; Kise, et al. 2022)].

The earliest molecular phylogenetic studies to employ complete mitochondrial genome sequences focused on relationships within Scleractinia (Medina, et al. 2006), Antipatharia (Brugler and France 2007), Hydrozoa (Kayal, et al. 2015), and across all of Cnidaria (Kayal and Lavrov 2008; Kayal, et al. 2013). More recently, cnidarian systematics has entered the phylogenomics age, with studies using data from hundreds (and sometimes thousands) of loci from transcriptome sequences (Chang, et al. 2015; Zapata, et al. 2015; Kayal, et al. 2018) and target-capture sequencing approaches (Quattrini, et al. 2018; Cowman, et al. 2020; Horowitz, et al. 2020; Quattrini, et al. 2020; Bentlage and Collins 2021; Glon, et al. 2021). We were able to tabulated 139 published cnidarian molecular phylogenetic studies focusing on major taxa (Table S1), but there are many more focused on smaller groups.

The accumulating body of phylogenetic evidence consistently recovers monophyletic Anthozoa, Hexacorallia, Octocorallia, Antipatharia, Ceriantharia, Zoantharia, Medusozoa, Staurozoa, Scyphozoa, Cubozoa, Hydrozoa, Endocnidozoa, and Myxozoa. However, the inferred phylogenetic relationships among and within these lineages differ in various studies. For example, many of the early cladistic and likelihood analyses of sequence data that included representatives of Staurozoa, Scyphozoa, Cubozoa, and Hydrozoa did not resolve the position of Staurozoa (Bridge, et al. 1995; Kim, et al. 1999; Collins 2002)(Fig. S1). Through analyses of morphology and 18S ribosomal RNA sequences, Marques and Collins (2004) found support for a clade consisting of Cubozoa and Staurozoa, with Scyphozoa as sister to this clade. Subsequent analyses of 28S ribosomal genes by Collins et al. (2006) supported Cubozoa and Scyphozoa as sister lineages, with Staurozoa as the sister group to this clade plus Hydrozoa. Kayal et al. (2013) found support in analyses of complete mitochondrial genome sequences for a clade that consisted of Staurozoa and Cubozoa, sister to a clade consisting of Hydrozoa and Scyphozoa. More recently, based on analyses of phylogenomic datasets, Zapata et al. (2015), Kayal et al. (2018), and Quattrini et al. (2020) all found support for Acraspeda, with Staurozoa as sister to the clade Rhopaliophora that unites Cubozoa and Scyphozoa (Fig. S1). No phylogenetic analysis published to date provides evidence to support recent taxa erected as part of a re-classification of acrasped cnidarians proposed by Straehler-Pohl and Jarms (2022a, b). Two main clades of hydrozoans, Trachylina and Hydroidolina, are consistently recovered as monophyletic, although studies using traditional Sanger sequencing markers have failed to recover relationships between the main lineages within Hydroidolina with sufficient support (e.g., Cartwright, et al. 2008; Collins, et al. 2008; Picciani, et al. 2018). Multiple studies have confirmed that most of the major groups of Hydroidolina -- Leptothecata, Siphonophorae, Capitata, and Aplanulata -- are monophyletic. However, Filifera has been found to be polyphyletic (Cartwright, et al. 2008; Collins, et al. 2008; Nawrocki, et al. 2010; Nawrocki, et al. 2013; Bentlage and Collins 2021). Historically, there has been little consistency in inferred relationships between higher-level groups within Hydroidolina. Recent phylogenetic analyses of Trachylina found congruent relationships among the major groups Limnomedusae, Trachymedusae, Narcomedusae, and Actinulida (Collins, et al. 2008; Bentlage, et al. 2018). Trachymedusae was found to be non-monophyletic, with one lineage derived from within Limnomedusae, and the rest of Trachymedusae paraphyletic with respect to Narcomedusae. To address part of this issue, Bentlage et al. (2018) revised Limnomedusae to include members of Geryonidae that were previously classified as Trachymedusae. As presently understood, Trachymedusae is still a paraphyletic assemblage that gave rise to a monophyletic Narcomedusae.

There has also been discordance in reconstructions of relationships within Anthozoa, perhaps with the most intriguing phylogenetic question being the placement of Ceriantharia. Analyses of 18S and 28S ribosomal RNA by Stampar et al. (2014) recovered Ceriantharia as sister to all other Hexacorallia. However, mitochondrial datasets placed Ceriantharia as sister to the rest of Anthozoa (Stampar, et al. 2014). Nuclear exon data from Zapata et al. (2015) and Kayal et al. (2018), ultraconserved element (UCE) data from Quattrini, et al. (2020) and studies of complete mitochondrial sequences from Stampar, et al. (2019) recovered the same result found in the ribosomal rRNA studies, with Ceriantharia as sister to the rest of Hexacorallia.

Phylogenomic studies (i.e. those with hundreds or thousands of loci sampled across the genome) have brought higher resolution to the cnidarian tree of life, but all of them lack taxonomic balance, and many omit key lineages (Fig. 1). For example, Zapata et al. (2015) included minimal ceriantharian and staurozoan data and did not include Myxozoa. Chang et al. (2015) added seven representatives of Myxozoa and a *Polypodium* transcriptome but lacked Staurozoa and Ceriantharia. Kayal et al. (2018) combined previous data sets and added five deeply sequenced transcriptomes from Staurozoa but included very few cerianthiarian and octocoral data and had as limited sampling within the most diverse clade of Hydrozoa, Hydroidolina. Kayal et al. (2018) and Zapata et al. (2015) resolved Aplanulata as the sister to a limited sampling of other Hydroidoloina. Bentlage and Collins (2021) tried to address this deficiency, using a bait capture approach focusing on Hydroidolina, and also recovered strong support for Aplanulata as the sister group to the remainder of Hydroidolina. As in past analyses, Bentlage and Collins (2021) found Filifera to be polyphyletic, but recovered support for a topology uniting Filifera I with Filifera II, as sister to Capitata, with these three taxa united in a clade sister to Leptothecata. This study also found support for Filifera III plus IV, as the closest relatives of Siphonophorae (Bentlage and Collins 2021). The UCE studies by Quattrini et al. (2018; 2020), which focused on Anthozoa, greatly increased the representation of that group, but by design included very few medusozoan taxa and only as outgroups. To comprehensively understand the evolutionary relationships among Cnidaria clades, it is essential to generate a phylogenetic tree that includes a comprehensive sampling across all major lineages.

**Fig. 1.**
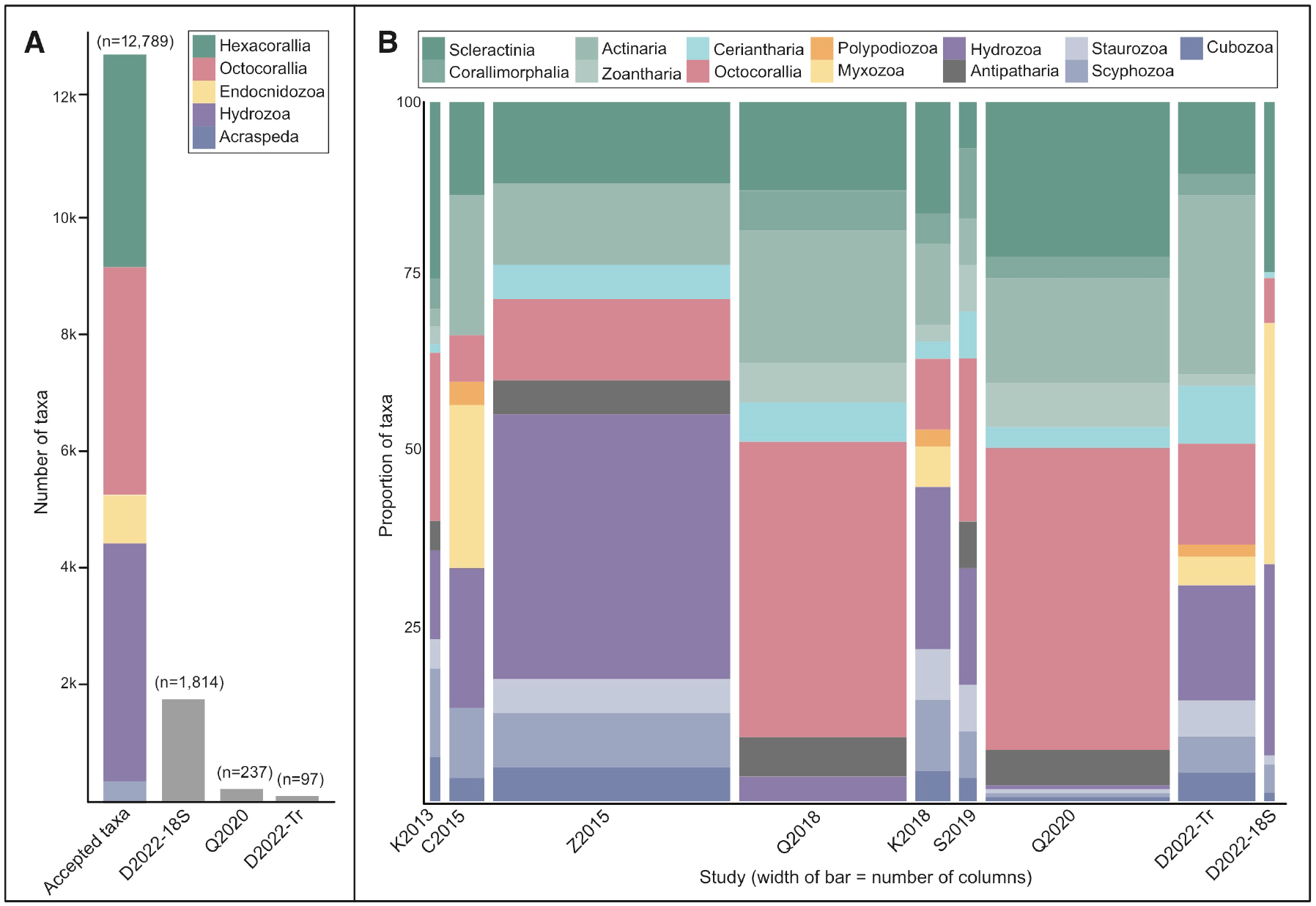
A graphical survey of cnidarian phylogenomic datasets over time, as compared to species richness across its major lineages. A) The colored bar represents the total number of cnidarian species described for four major taxonomic groups. The number of accepted taxa is based on the World Register of Marine Species database as of September 2022. The height of each grey bar represents the number of species from the total described included in the corresponding study. B) The height of each colored section represents the proportion of a particular taxonomic group included in the study. The width of the bar represents the number of nucleotide or amino acid columns in the dataset. For panels A and B, the studies are abbreviated with the first letter of the first author’s surname and the publication year (e.g., K2013 represents Kayal et al. 2013). D2022-18S and D2022-Tr indicate the 18S and transcriptomic datasets generated in this study, respectively. The other studies included are Chang et al. 2015, Zapata et al. 2015, Quattrini et al. 2018, Kayal et al. 2018, Stampar et al. 2019, and Quattrini et al. 2020.

Here, we combine 26 de novo transcriptome datasets and previously published transcriptome and gene model datasets to increase taxon sampling for underrepresented clades and improve the balance of taxon sampling across Cnidaria. We use the topology resulting from phylogenomic analyses of our transcriptome data to constrain a phylogenetic analysis of more than 1,800 small subunit ribosomal DNA (18S rDNA) sequences. Our resulting phylogenies and new transcriptomic data provide a solid framework for present understanding of the evolutionary history of Cnidaria, and for guiding future research on the phylogenetics of Cnidaria.

## Methods

### Reproducibility and transparency statement

Custom scripts, command lines, and data used in these analyses, including transcriptomes, and alignment and tree files, are available at our GitHub repository (https://github.com/josephryan/DeBiasse_cnidophylogenomics) and Dryad (https://doi.org/10.6071/M3K39S). To maximize transparency and minimize confirmation bias, we planned analyses *a priori* using phylotocol (DeBiasse and Ryan 2019) and posted this original document and any subsequent changes to our GitHub repository.

### Sample collection and data generation

We generated new transcriptome data for 26 anthozoans (Table S2). Following the methods described in Pelosi et al. 2022, we generated RNA-Seq data for *Eunicea flexuosa, Eunicea tourneforti*, and *Muricea muricata* from individuals collected at Media Luna reef, Puerto Rico, in April 2013. Following the methods described in Klompen et al. (2020), we generated RNA-Seq data for *Isarachnanthus maderensis* and *Botruanthus mexicanus* from individuals collected in Sisal, Yucatan, Mexico in August 2018 and for *Arachnanthus* sp. from an individual collected in Barra del Chuy, Rocha, Uruguay in March 2019. Following the methods described in Macrander et al. (2015, 2016) we generated RNA-Seq data for the following species: *Bartholomea annulata* and *Bunodeopsis antilliensis* collected from Grass Key, Marathon, Florida, in June 2013; *Lebrunia neglecta* collected in the Florida Keys in June 2103; *Bunodosoma cavernatum* collected from Galveston, Texas, in October 2013; *Diadumene leucolena, Diadumene lineata*, and *Haloclava producta* collected from Woods Hole, Maine in May 2013; *Stomphia coccinea* collected from Friday Harbor, Washington in September 2013; *Condylactis gigantea* and *Entacmaea quadricolor* purchased from a PetCo retail store in Columbus, Ohio (collection location unknown) in 2012; *Triactis producta* (collected in Indonesia), *Calliactis polypus* (collection location unknown), and *Macrodactyla doreensis* (collection location unknown), purchased from LiveAquaria.com in September 2013; *Actinia equina* collected from Antrim, Northern Ireland, UK, in August 2013; *Andwakia discipulorum* collected from Kana’ohe Bay, Oahu, Hawaii in 2012; *Epiactis prolifera* collected from Bodega Bay, California in 2012; *Cylista elegans* collected from Strangford Lough, Northern Ireland in 2013; *Metridium senile* collected from Darling Marine Lab, Damariscotta, Maine in 2013. The *Leptogorgia* sp. sample was collected and RNA was extracted at the University of Florida Whitney Lab for Marine Bioscience in Flagler County, FL in February 2017. Adult *Ceriantheopsis americana* individuals were collected from Cedar Key in Levy County, FL in March 2018. In April 2018, a larva resulting from the spawning of the adults was collected and RNA was extracted following the protocol in Babonis et al. 2016. RNA-Seq libraries for *Leptogorgia* sp. and *C. americana* were prepared and 150bp paired end reads were sequenced on the Illumina HiSeq 3000 platform at the University of Florida Interdisciplinary Center for Biotechnology Research.

### Transcriptome assembly and processing

We trimmed FASTQ sequences and assembled transcriptomes using the cutadapt option in Trinity v2.8.5 (Grabherr, et al. 2011). We applied the --include_supertranscripts parameter to generate superTranscripts as part of each Trinity run. SuperTranscripts provide a single all-inclusive transcript for genes with multiple isoforms (Davidson, et al. 2017). We translated the superTranscripts into amino acid sequences in TransDecoder v5.0.2 (github.com/TransDecoder). We set the TransDecoder ‘-m’ flag (minimum length of open reading frame) to 50 and used the results from BLASTP (McGinnis and Madden 2004) searches to inform the final TransDecoder prediction step. We filtered potential contaminants in these translated sequences by removing sequences where the top BLASTP hit to v0.02 of the curated alien_index database (http://ryanlab.whitney.ufl.edu/downloads/alien_index/) was to a non-metazoan sequence using Alien Index v2.1 (https://github.com/josephryan/alien_index). We assessed the completeness of each transcriptome by searching against the eukaryote database in BUSCO v2 (Simão, et al. 2015) as implemented in gVolante v1.2.0 (Nishimura, et al. 2017).

### Phylogenomic matrix construction and phylogeny estimation

Our original dataset consisted of 26 new anthozoan transcriptomes, four cnidarian transcriptomes that we assembled from data publicly available on the NCBI Short Read Archive, 75 previously assembled and published transcriptomes, and seven previously published amino acid gene model data sets (112 sequences total, Table S2). The dataset included 104 cnidarians and eight outgroup taxa (Table S2). We used diamond v0.9.22.123 (Buchfink, et al. 2015) to perform reciprocal best BLAST searches and generated FASTA files of orthologous sequences (i.e., orthogroups) in OrthoFinder v2.2.3 (Emms and Kelly 2019) using all 112 sequences as input.

We filtered the orthogroups inferred by OrthoFinder as follows: using an automated script, sequences within each orthogroup were aligned using MAFFT v7.309 (Katoh and Standley 2013), and in the multicore version of IQ-TREE v1.5.5 (Nguyen et al. 2015), a maximum likelihood (ML) tree for each alignment that had no more than 50% sequence gaps was estimated. Only the orthogroup trees that had at least 50% of the total taxa and no more than eight paraphyletic duplicates per species were retained (there were no limits on the number of duplicates if they were monophyletic). In PhyloTreePruner v1.0 (Kocot, et al. 2013), all but one sequence in taxa with monophyletic duplicates (e.g., paralogs) were removed, which produced a set of orthologous loci with one sequence per species in at least 50% of our taxa. Because the initial run of the orthogroup filtering pipeline produced a small number of single-copy loci, we removed five cnidarian species that had a high number of duplicates per core gene in BUSCO analyses (Table S2). We removed *Heteractis crispa* because it clustered with the outgroup taxa in preliminary trees and subsequent BLAST analyses suggested the *H. crispa* transcriptome was substantially contaminated with vertebrate sequences. *Muricea muricata* was also removed due to suspected non-target cnidarian contamination. After removing these seven taxa, we reran OrthoFinder and the filtering pipeline with the parameters described above for the 105 species remaining.

We concatenated alignments of all of the single-copy orthologs for the remaining 105 species using the fasta2phylomatrix utility v0.02 (github.com/josephryan/JFR-PerlModules) and aligned these sequences with MAFFT v7.309. We used this matrix for downstream analyses. This dataset did not involve any column trimming, as it has been shown that current methods for filtering multiple sequence alignments lead to suboptimal alignments (Tan, et al. 2015). Nevertheless, to test if removing divergent and ambiguously aligned columns affected our results, we generated a trimmed version of this matrix with Gblocks v0.91b (Castresana 2000) using dynamic parameters generated by Gblockswrapper v0.03 (https://goo.gl/fDjan6).

We used both matrices (trimmed and untrimmed) and multiple models (partitioned and unpartitioned) to estimate four ML phylogenies in IQ-TREE v1.5.5 (Nguyen, et al. 2015). In the first and second analyses, we used the IQ-TREE parameter ‘-m TEST’ to determine site-homogeneous models of amino acid substitution for each gene partition applied to the (i) untrimmed and (ii) trimmed data matrices. In the third and fourth analyses, we used the C60 model in IQ-TREE, which accounts for across-site compositional heterogeneity in equilibrium frequencies, applied to the (iii) untrimmed, unpartitioned data matrix and the (iv) untrimmed, partitioned data matrix. Support values for all phylogenies were determined from 1000 bootstrap replicates.

### Small subunit (18S) ribosomal DNA matrix construction

We ran the following search at GenBank (NT) on July 8, 2020: ((Cnidaria[ORGN] AND (18S OR “small subunit ribosomal”)) BUTNOT Nematostella[ORGN]) OR AF254382. We downloaded these results in GenBank format. To these 13,717 sequences, we added accessions AF254382 (*Nematostella vectensis*) and AF052581 (*Renilla reniformis*). We used a custom script (get_18S_fasta_from_genbank.pl) to convert GenBank format to FASTA and remove the following sequences: (1) all duplicates of a species except for the longest, (2) AY935208 (*Aurelia* sp.), (3) sequences shorter than 1000 nucleotides, (4) sequences that include the patterns ‘environ,’ ‘parasite,’ or ‘proliferative’ in their definition line, (5) sequences that did not include a class designation, and (6) sequences from taxa that include species *affinis* (abbreviated sp. or cf.) unless those sequences were the only representative of a genus.

The following changes were made based on prior knowledge: (1) *Virgularia gustaviana* was removed as it is erroneously annotated (clearly a ceriantharian) in GenBank, (2) *Carybdea marsupialis* was renamed *Alatinidae* indet., (3) *Alatina philippina* was removed as it was shown to be the same as *Alatina morandinii* (Straehler-Pohl and Toshino 2015), (4) *Darwin* sp. was renamed to *Gerongia rifkinae*, and (5) accession AF099104 (*Craterolophus convolvulus*) is a contaminant of *Haliclystus* so it was replaced with AY845344. After running an initial tree, we identified one long-branched clade of octocoral sequences that contained *Junceella aquamata* (AY962535), *Junceella fragilis* (AY962533), and *Subergorgia ornata* (AY962537), which fell within Hexacorallia instead of Octocorallia. We determined that these three sequences were likely contaminants and removed them based on the following criteria: (1) all were from the same NCBI PopSet (accession=63148780), (2) the top BLAST hits of each of these were to other sequences from this PopSet including bivalves and crustaceans, (3) *Junceella* and *Subergorgia*, which were sister in our prelim tree, are distant genera in Quattrini et al. (2020), and (4) there is no obvious voucher available for these sequences.

We used ssu-align v0.1.1 (Nawrocki and Farm 2010) with default settings to align the 18S sequences. We used ssu-mask v0.01 from the same package with default settings to remove columns that likely include a significant number of misaligned nucleotides as recommended in the ssu-align manual. We used esl-format v0.43 from the Easel sequence library (https://github.com/EddyRivasLab/easel) to convert stockholm formatted alignments to FASTA format.

### Small subunit (18S) ribosomal DNA phylogeny estimation

We constructed a constraint tree based on the transcriptome-based phylogeny. We pruned *Cerianthopsis americana* from the constraint tree because there has been confusion regarding the distinction between this taxon and *Pachycerianthus borealis* (Klompen, et al. 2020) that complicates downstream interpretations. Additional sequences were pruned if a corresponding sequence did not exist in the 18S dataset. In total, there were 23 sequences pruned from the constraint tree (Table S2). We also collapsed the three cubozoan species into a polytomy to reflect discrepancies that we encountered among the phylogenomic trees (discussed in Results).

We next generated a phylogeny using IQ-TREE multicore v1.6.12 (Nguyen, et al. 2015) applying the TN model (Tamura and Nei 1993) with a 4-class gamma distributed rate heterogeneity (G4). The final dataset, the accessions of all sequences, the constraint tree, the final tree, and the scripts used to create the dataset and constraint tree, are posted to our GitHub repository (URL above) and the transcriptomes are available at Dryad (https://doi.org/10.6071/M3K39S).

## Results

### Phylogenomic analyses

We generated new transcriptome data for 26 anthozoan species: 18 Actiniaria (Hexacorallia), four Ceriantharia (Hexacorallia), and four Octocorallia. To these we added 86 publicly available transcriptome and gene model datasets. After performing quality control, we removed seven transcriptomes from the analyses because we detected high gene duplication or contamination, leaving 97 cnidarian species and eight non-cnidarian outgroups in our analyses (Table S2).

We assigned 3,670,777 of 4,892,912 sequences (75%) to orthogroups. We retained 4,117 orthology groups that had at least 53 of 105 species present and no more than eight duplicates per species. After removing within-species duplicates that were determined to be paralogs or isoforms, we were left with 748 single-copy orthogroups. After alignment, the resulting matrix consisted of 449,935 amino acid columns. We also generated a trimmed version of this matrix containing the 123,051 most unambiguously aligned amino acid columns.

Phylogenies inferred using the trimmed and untrimmed matrices under partition-specific, site-homogeneous models in IQ-TREE (i & ii) were largely concordant in terms of support values and topology with a few exceptions (Fig. 2 and S2). In the two phylogenies estimated under the C60 model (iii & iv), Staurozoa was sister to Hydrozoa with low support (Fig. S3 and S4). In the phylogeny estimated from the untrimmed and unpartitioned matrix under the C60 model (iii, Fig. S3), the myxozoan taxa *Thelohanellus kitauei* and *Myxobolus cerebralis* were sister, while in the other three phylogenies we recovered *T. kitauei* as sister to *Myxobolus pendula* (Fig. 2, S2, S4). In the phylogeny estimated from the untrimmed matrix under the partition-specific site-homogeneous model (ii, Fig. 2), the cubozoan taxa *Alatina alata* and *Chironex fleckeri* were sister, while in the other three phylogenies, *A. alata* was sister to *Tripedalia cystophora* (Fig. S2-S4). All four phylogenies we estimated placed Ceriantharia sister to other Hexacoralia with full bootstrap support (Fig. 2, S2-S4). In all four phylogenies, Enthemonae and Anenthemonae were monophyletic, but within Enthemonae, *Haloclava producta* was nested within Metridioidea, making Metridioidea and Actinioidea nonmonophyletic (Fig. 2, S2-4).

**Fig. 2.**
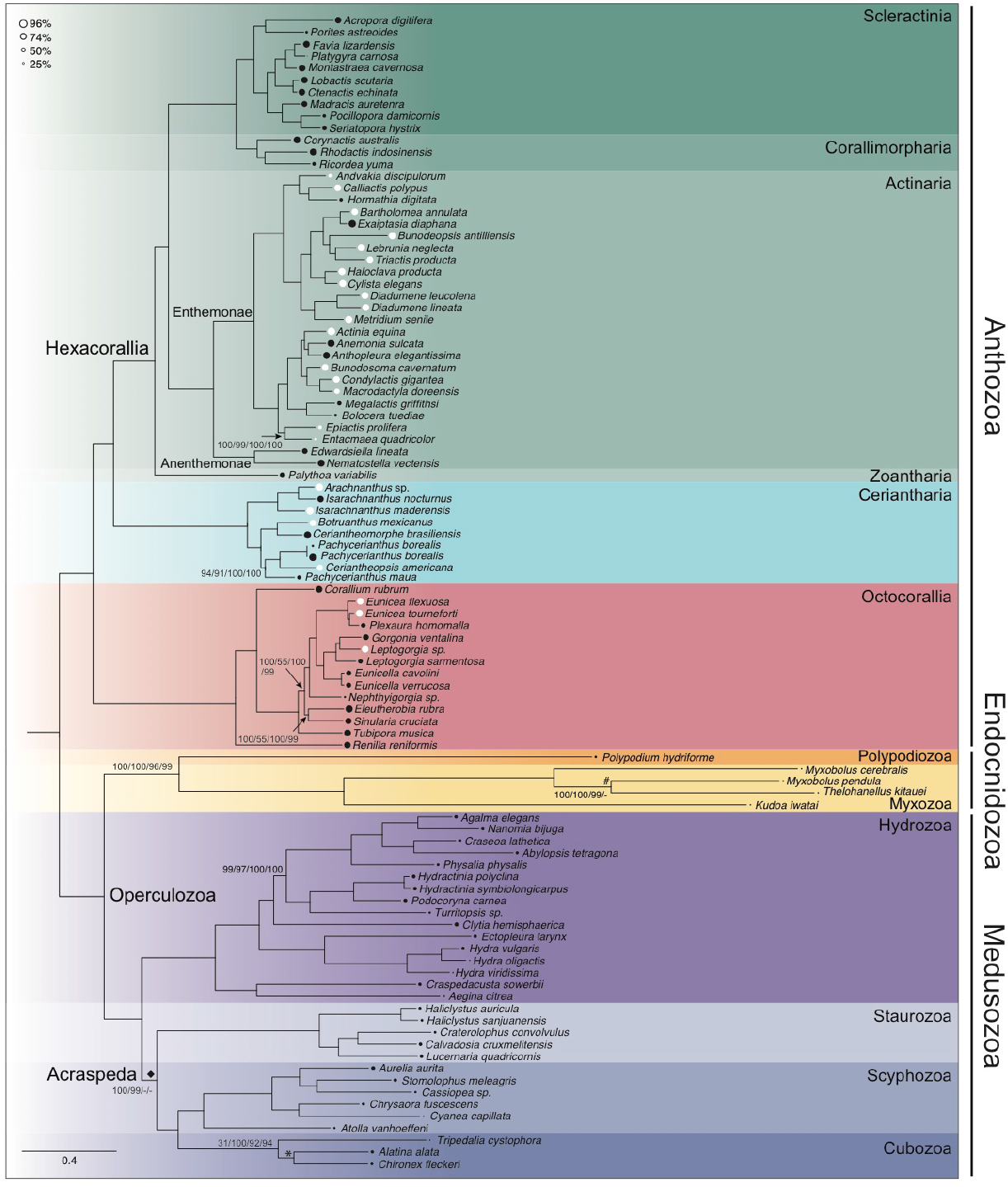
Maximum-likelihood phylogeny of Cnidaria. estimated from an untrimmed, concatenated matrix of 748 ortholog alignments analyzed under a site homogeneous partitioned model (rooted based on the outgroup, which is not shown). Circles at the branch tips are proportional to the occupancy of that taxon in the data matrix, with black circles indicating previously published data and white circles indicating data generated for this study. Occupancy for previously published Myxozoa, *Cyanea capillata, Tripedalia cystophora, Platygyra carnosa* sequences was below 15% (Table S2), making circles for these taxa very small and appearing to be missing. Bootstrap values are indicated at nodes if support is less than 100% for any of the analyses (partition-specific site-homogeneous models run on untrimmed matrix listed first, partition-specific site-homogeneous models run on trimmed matrix listed second, C60 analysis listed third, and C60 partitioned analysis listed fourth). The hash indicates a conflicting relationship where *T. kitauei* is sister to *M. cerebralis* in all other analyses (Fig. S2-S4). The diamond indicates a conflicting relationship in the two phylogenies estimated under the C60 model where Staurozoa is sister to Hydrozoa (Fig. S3-S4). The asterisk indicates a conflicting relationship where *A. alata* is sister to *T. cystophora* in the three other phylogenies (Fig. S2-S4).

### Small subunit (18S) ribosomal DNA phylogeny

We downloaded 13,717 small subunit ribosomal (18S) gene sequences from GenBank. After choosing a single representative from sets of sequences from the same species name (retaining the longest sequence of a set of duplicates), and removing sequences deemed problematic for various reasons (see methods), we built a matrix consisting of 1,814 sequences (15 Ceriantharia, 442 non-ceriantharian Hexacorallia, 117 Octocorallia, 496 Hydrozoa, 22 Cubozoa, 24 Staurozoa, 73 Scyphozoa, and 625 Endocnidozoa) representing 702 genera. The resulting alignment of these sequences consisted of 1,324 columns (959 parsimony-informative). We constructed a constraint tree based on the untrimmed, partitioned site homogeneous analysis above (Fig. 2). This tree included 75 taxa that were both in our phylogenomic dataset and our 18S analysis. Cubozoa was collapsed to a polytomy in our constraint tree to reflect the disagreement among phylogenetic analyses. We then applied this constraint tree in a maximum-likelihood analysis.

The resulting analysis produced a cnidarian tree that included roughly 15% of all described cnidarian species (Fig. 3). Of the 229 genera with more than a single representative, 47 (21%) were identified as monophyletic in our tree.

**Fig. 3.**
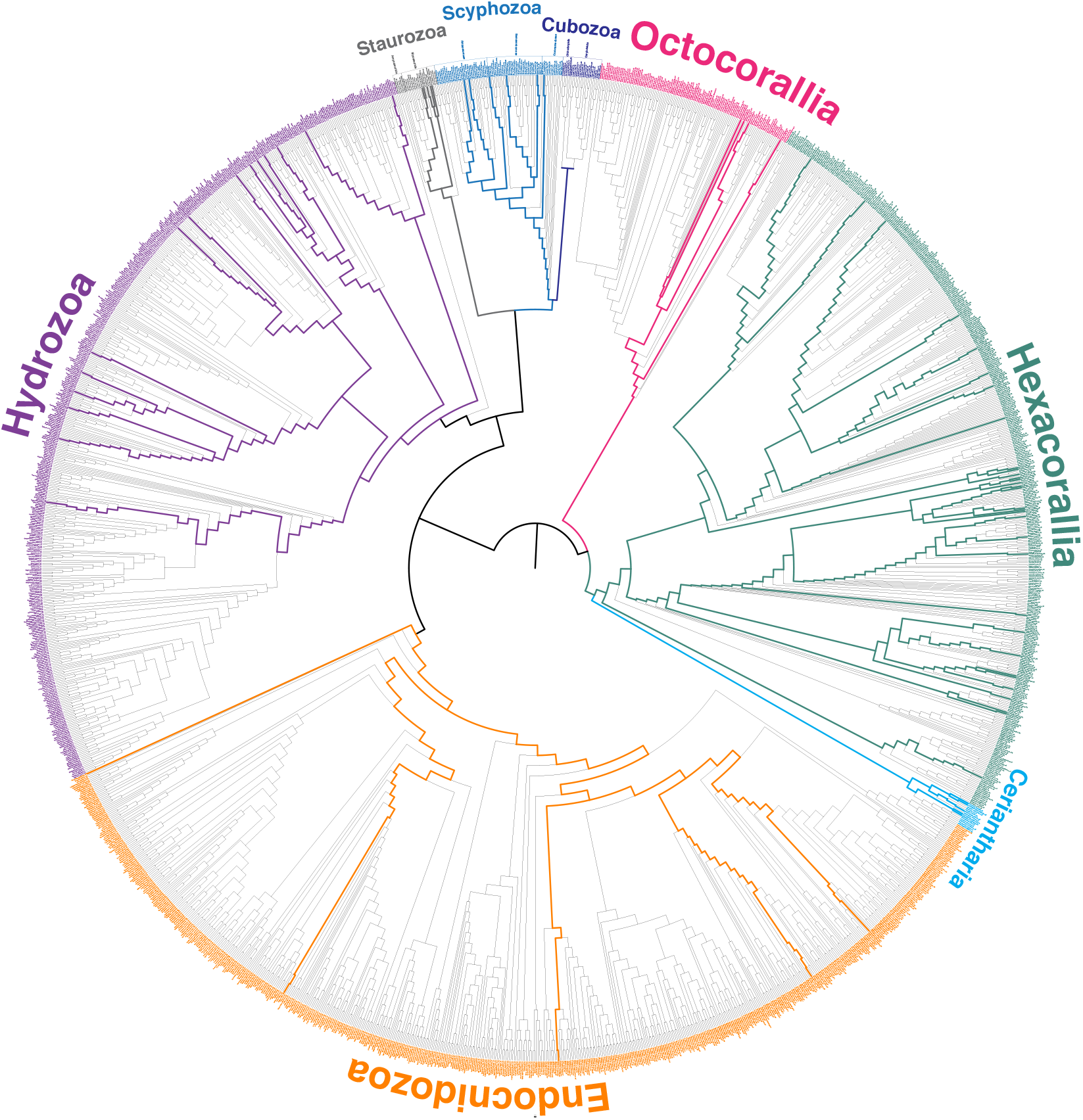
Maximum likelihood phylogeny estimated using small subunit ribosomal (18S) gene sequences from 1,1814 cnidarian species with higher-level relationships constrained according to the relationships in the multi-locus, untrimmed, partitioned, site homogeneous analysis (Fig 2). Bolded branches indicate the constraint tree applied, with colors representing higher-level taxonomic groups as in Fig. 1 and 2.

## Discussion

Cnidaria is an ancient lineage that encompasses a wide range of phenotypic and genomic diversity. Research interests in cnidarian organisms are extensive and taxonomically broad, presenting an excellent opportunity to study a wide variety of biological processes. However, an accurate and extensive phylogenetic framework is necessary to promote and contextualize such research. Existing phylogenies encompass less than 2% of species (Fig. 1A) and most, if not all, are taxonomically skewed relative to the actual representation of major cnidarian groups (Fig. 1B). In addition, there are conflicting relationships that emerge from these previous studies, which is not surprising given that they differ in taxon sampling and fail to represent the diversity of the group. Here, we apply a hybrid approach to present phylogenetic relationships for nearly 15% of cnidarian species. Building a reliable phylogeny for any group, particularly one as diverse and species rich as Cnidaria, requires the cumulative efforts of researchers working to improve inference methods and taxonomic and genomic sampling. In comparing results across studies and incorporating previously applied sequences with newly acquired data, support for phylogenetic relationships becomes stronger.

### Anthozoa

#### Hexacorallia

The main phylogeny generated here (Fig. 2) recovers all included ordinal lineages as monophyletic and concurs with other recent phylogenomic studies (Zapata, et al. 2015; Kayal, et al. 2018; Quattrini, et al. 2018; Quattrini, et al. 2020) in finding Ceriantharia as the sister to all other hexacorals as well as recovering Scleractinia and Corallimorpharia as sister taxa. We also find Zoantharia as the sister to a clade that includes Scleractinia, Corallimorpharia, and Actiniaria, but the absence of Antipatharia in our phylogenomic analyses limits comparison beyond that broad frame. In the much more densely sampled 18S tree, Zoantharia is non-monophyletic with a sample labeled as *Zoanthus* falling out among the Antipatharia; we suspect that this is a contaminant or mislabeled sequence rather than evidence for zoantharian polyphyly. Ordinal-level relationships in our 18S tree contravene previous phylogenomic interpretations, with Antipatharia as the sister to the Actiniaria-Scleractinia-Corallimorpharia clade, rather than being sister to Scleractinia and Corallimorpharia as in target-enrichment (Quattrini, et al. 2018; Quattrini, et al. 2020) and transcriptome-based analyses (Zapata, et al. 2015). Our reconstruction of ordinal phylogeny conflicts with any inference of homology for attributes related to skeletonization of hexacorals, as it disassociates Scleractinia and Antipatharia.

#### Ceriantharia

Among anthozoan lineages, Ceriantharia has been the most challenging to interpret phylogenetically. Instability in the resolution of Ceriantharia confounds attempts to understand two key aspects of ceriantharian biology. Ceriantharians produce spirocysts and ptychocysts in addition to nematocysts, with these additional kinds of capsules having different structural properties and functions (Mariscal 1984). Ptychocysts are unique to Ceriantharia (Mariscal, et al. 1977). The inferred relationship among these secretory products depends on the topology of the anthozoan tree (Reft and Daly 2012) and is worth investigating, given the functional importance and unparalleled complexity of these microscopic machines. Similarly, the medusiform, long-lived larval stage of some ceriantharians is complicated to interpret if the phylogenetic position of Ceriantharia is not well resolved.

Here we applied the greatest diversity and number of ceriantharian species to date (Fig. 1B, Fig. 2), and recovered this historically labile group as sister to other Hexacorallia in all of our analyses (Fig. 2, S2–4). This result corroborates the results of previous phylogenomic studies (Zapata, et al. 2015; Kayal, et al. 2018; Quattrini, et al. 2018; Quattrini, et al. 2020) and solidifies the placement of this clade. With this result in hand, we can gain insight from prior studies that recovered a conflicting result. Growing evidence that these relationships are outliers lends support to the hypothesis that the topology resolved in Stampar et al. (2019) is the result of unique and remarkably slow rates of mitochondrial genome evolution (Shearer, et al. 2002; Brugler 2004; Hellberg 2006), a trend echoed across most anthozoan groups. In the phylogenomic tree, Ceriantharia contains a monophyletic Penicillaria and Spirularia. However, within Spirularia, the two Cerianthidae species are not monophyletic, as *Pachycerianthus borealis* (Cerianthidae) is sister to a clade that contains *Botruanthus benedeni* (Botrucnidiferidae) and *Ceriantheomorphe brasiliensis* (Cerianthidae). We also find the genus *Isarachnanthus* non-monophylteic as *I. nocturnus* is sister to *Archnanthus* sp.

Our constrained 18S tree presents further conflicts between taxonomy and phylogeny. Here the genus *Pachycerianthus* is non-monophyletic as a clade of six *Pachycerianthus* species is sister to the two *Isarachnanthus* (Penicillaria) species. In addition, the two Botrucnidiferidae taxa (*Botruanthus benedeni* and *Botrucnidifer* sp.) and the genus *Cerianthus* are non-monophyletic. These problems echo results of previous molecular phylogenetic relationships among Ceriantharia, which generally find conflict between taxonomic groups and phylogenetic results (e.g., Stampar, et al. 2012; Stampar, et al. 2014; Forero Mejia, et al. 2020). The disconnect between taxonomy and phylogeny in both the phylogenomic and 18S data are paralleled in recent discoveries of significant plasticity in morphology within the life history of a species and persistent taxonomic confusion, even at a narrow geographic scale (reviewed in Stampar, et al. 2016).

#### Actiniaria

Our study introduces 18 new actiniarian transcriptomes. Although we found monophyly for Actiniaria as a whole, internal relationships differed significantly from previous studies. For example, in our 18S tree, Enthemonae and Anenthemonae are non-monophyletic, in contrast to previous studies (Rodríguez, et al. 2014; Gusmão, et al. 2020), with the difference stemming from the placement of the actinostolideans *Hormosoma* and *Anthosactis* and the actinernoideans *Actinernus, Isactinernus*, and *Synactinernus* at the base of the actiniarian tree rather than within Enthemonae and Anenthemonae, respectively. This novel topology recalls historical groupings of these taxa within “Mesomyaria,” (see Carlgren 1949; Rodríguez, et al. 2014) and if confirmed, would significantly change the inferred history and homology of marginal musculature in Actiniaria. Based on the 18S tree, the marginal sphincter would be inferred to be present and mesogleal at the root of Actiniaria, with subsequent losses and multiple shifts to become concentrated and embedded in the endoderm. Internal relationships in Anenthemonae are unconventional, including a non-monophyletic Actinioidea and Metridiodea. The phylogenomic results are less controversial, finding monophyly for Anenthemonae, Enthemonae, Actinioidea, and Metridioidea. However, the phylogenomic data lacks mesomyarian taxa, which typically branch near the base of the actiniarian tree. Furthermore, the taxon sampling in the phylogenomic dataset is almost an order of magnitude lower for Actinoidea and Metridioidea than that of the 18S tree.

#### Scleractinia and Corallimorpharia

Reconciling molecular phylogeny with morphology in Scleractinia has been a long term, persistent, and challenging task (McMillan, et al. 1991; Fukami, et al. 2004; Fukami, et al. 2008; Kitahara, et al. 2010), with Huang et al. (2011) coining the tongue-in-cheek term “Bigmessidae” to describe this group. Some of the earliest molecular phylogenies based on mitochondrial rRNA genes resulted in two major scleractinian groups that conflict with morphologically defined subordinal classifications (McMillan, et al. 1991; Romano and Palumbi 1996, 1997; Romano and Cairns 2000; Chen, et al. 2002). One group, called the robust corals, contains platelike and massive taxa with thick, heavily calcified skeletons. The second group, called the complex corals, contains corals with more porous, less calcified skeletal walls. We recovered these robust and complex groups (Fig. 2, S2-4) as have recent multilocus studies (e.g., Lin, et al. 2016). An important area of future research attention is the position of deep-sea aposymbiotic taxa, which fall out as the earliest branching scleractinians in studies based on mitogenomes and rRNA (Barbeitos, et al. 2010; Kitahara, et al. 2010; Stolarski, et al. 2011; Seiblitz, et al. 2020), but have been absent or underrepresented in recent phylogenomic studies.

#### Introducing Coralliforminae (Corallimorpharia + Scleractinia)

Using whole mitochondrial genomes, Medina et al. (2006) found Corallimorpharia sister to the complex corals, rendering Scleractinia non-monophyletic. Based on these relationships, Medina et al. (2006) resurfaced the “naked coral hypothesis,” suggesting the soft-bodied corallimorpharians evolved from a scleractinian ancestor and subsequently lost the stony skeleton trait during historical periods of increased CO_2_ concentrations in the marine environment. Subsequent studies using a range of loci (nuclear rRNA: Fukami, et al. (2008); whole mitochondrial genomes: Kayal, et al. (2013); Lin, et al. (2014); Seiblitz, et al. (2020); nuclear protein coding: Lin, et al. (2016); Kayal, et al. (2018); UCEs: Quattrini, et al. (2018); Quattrini, et al. (2020)) have found Scleractinia to be monophyletic. Like these studies, we recovered a monophyletic Scleractinia across all four phylogenetic analyses (Fig. 2, S2–4), demonstrating this relationship is robust to model choice and refuting the naked coral topology, which was likely a result of saturation in the mitochondrial protein sequences, long-branch attraction, and/or model violations (Kitahara, et al. 2014). We propose the name Coralliformes to represent the clade that unites Corallimorpharia and Scleractinia.

#### Octocorallia

In all four of our phylogenetic analyses, we found Octocorallia sister to Hexacorallia (Fig. 2, S2–4), as have recent phylogenomic studies (Quattrini, et al. 2018; Quattrini, et al. 2020), demonstrating the stable position of octocorals in the cnidarian tree. However, ordinal and familial-level taxonomy has been, and continues to be, uncertain (reviewed in McFadden, et al. 2021). Recently, Quattrini et al. (2018; 2020) made major strides in octocoral phylogenomics, increasing the number of taxa (94 Alcyonacea, 2 Heliporacea, 7 Pennatulcea) and loci (933 UCE loci, 278,819bp). Nevertheless, some discrepancies between taxonomy and molecular phylogeny remain. For example, while Helioporacea and Pennatulacea were monophyletic, each was nested within Alcyonacea (Quattrini, et al. 2020). Lower-level taxonomy in octocorals also remains unresolved. In our transcriptomic tree, we find the genus *Leptogorgia* non-monophyletic and the suborder Alcyoniina non-monophyletic as *Nephthyigorgia* is sister to clade of *Holaxonia*, although this relationship may be unreliable because *Nephthyigorgia* has very low matrix occupancy (30%, Table S2). Future studies would benefit from increased evenness in taxonomic sampling. For example, Alcyonacea is overrepresented in recent phylogenomic studies (Quattrini, et al. 2020). Here, we have only one representative of Pennatulacea (*Renilla reniformis*) and had to remove the only representative of Helioporacea (*Heliopora coerulea*) due to high numbers of transcript copies and/or paralogs (Table S2). In light of the extensive UCE-based phylogeny (Quattrini, et al. 2018; Quattrini, et al. 2020) and the long-realized unreliability of 18S for constructing octocoral relationships due to its lack of sufficient variation among octocoral taxa (McFadden, et al. 2010), we do not go into extensive details of the octocoral relationships in our 18S tree.

### Endocnidozoa

We find support for Endocnidozoa, recovering Myxozoa sister to Polypodiozoa (Fig. 2, S2–4), concordant with Chang et al. (2015) and Kayal et al. (2018). Endocnidozoa is another group that would benefit from improved taxon sampling in cnidarian phylogenetic studies. Here, we included data for the same five species included in Kayal et al. (2018) with low matrix occupancy (5-35%, Table S2). Chang et al. (2015) included data for 8 species in a study specifically designed to test the placement of this group within Cnidaria. Neither Zapata et al. (2015) nor Quattrini et al. (2020) included endocnidozoan taxa. Across our four analyses, we found two topologies, both with paraphyletic *Myxobolus*. In the site-heterogeneous unpartitioned analyses, *Thelohanellus kitauei* and *M. cerebralis* formed a clade (Fig. S3) and in all other analyses, *T. kitauei* formed a clade with *M. pendula* (Fig. 2, S2, S4).

Despite variation in the relationships within Myxozoa, we found support for the larger clade, recovering Myxozoa sister to Polypodiozoa across all four analyses (Fig. 2, S2–4), concordant with Chang et al. (2015) and Kayal et al. (2018). Another relationship consistent in past studies (Chang et al. 2015, Kayal et al. 2018) and in all four analyses performed here is the sister relationship of Endocnidozoa and Medusozoa (Fig. 2, S2–4, and see next section).

### Introducing Operculozoa (Medusozoa + Endocnidozoa)

As stated above, a relationship consistent in past studies (Chang et al. 2015, Kayal et al. 2018) and in all four analyses performed here is the sister relationship of Endocnidozoa and Medusozoa (Fig. 2, S2–4). Based on these results, and noting that nematocysts of Medusozoa and Polypodiozoa (see Raikova and Raikova 2016) and polar capsules of Myxozoa all possess an operculum (Reft and Daly 2012), we propose the name Operculozoa for the clade uniting Endocnidozoa and Medusozoa.

### Medusozoa

Whereas our study has served to solidify relationships within Anthozoa, our results highlight the need for more taxon sampling within Medusozoa. Relationships of the major medusozoan lineages have been increasingly refined in recent decades (Bridge, et al. 1995; Kim, et al. 1999; Collins 2002; Marques and Collins 2004; Collins, Bentlage, et al. 2006; Kayal, et al. 2015), with an emerging consensus that Medusozoa consists of two major lineages, Acraspeda and Hydrozoa (Kayal et al. 2018). In our analyses that employed 20-states empirical amino acid substitution matrices (i & ii), we recovered a monophyletic Acraspeda with Staurozoa as sister to the clade containing Scyphozoa and Cubozoa, matching the relationships found by four recent studies (Chang et al. 2015, Zapata et al. 2015, Kayal 2018, Quattrini et al. 2020). However, we recovered an intriguing, albeit weakly supported, finding when applying a site-heterogeneous model, in which Hydrozoa and Staurozoa formed a clade sister to a clade containing Scyphozoa and Cubozoa (Fig. S1, S3, S4), a result also recovered by Miranda et al. (2016) analyzing concatenated mitochondrial (COI, 16S) and nuclear (ITS, 18S, 28S) loci under a Bayesian framework. None of our results corroborate the ribosomal RNA-based hypothesis that Staurozoa is sister to the remaining medusozoans (Collins, Schuchert, et al. 2006; Picciani, et al. 2018).

Site-heterogeneous models that partition data at the level of sites (as opposed to at the level of genes) have been suggested to alleviate problems of long-branch attraction (Lartillot and Philippe 2004; Le and Gascuel 2008). It is possible that our recovery of a non-monophyletic Acraspeda was due to applying a site-heterogeneous approach. However, Kayal et al. (2018) applied the site-heterogeneous CAT model in PhyloBayes (Lartillot, et al. 2009) using largely the same medusozoan taxa as we used here and recovered a monophyletic Acraspeda. These conflicting relationships within Medusozoa may be a result of a number of factors including: (1) increased anthozoan sampling in our study, (2) increased gene sampling in our study, or (3) differences between the C60 model implemented in IQ-TREE and the CAT model implemented in PhyloBayes. Nevertheless, taxon sampling within Medusozoa is poor in all phylogenomic analyses to date and a true understanding of medusozoan relationships will require substantial increase in data from this group (Fig. 1).

### Cubozoa

In addition to conflicting relationships between the major medusozoan lineages, we also find unstable relationships within Cubozoa across our analyses. We included three cubozoans in our analyses: *Chironex fleckeri, Tripedalia cystophora*, and *Alatina alata*. In our untrimmed site-homogeneous analysis, we recovered *Tripedalia cystophora* as sister to a clade containing *Chironex fleckeri* and *Alatina alata* (Fig. 2). We also recovered this relationship in our 18S phylogeny, which did not constrain relationships within Cubozoa. These results conflict with the rest of our analyses as well as with Bentlage, et al. (2010) and Kayal et al. (2018), which recovered *C. fleckeri* (Chirodropidae) as sister to a clade containing *A. alata* (Alatinidae) and *T. cystophora* (Tripedaliidae). This inconsistency is almost certainly due to the poor taxon sampling (3 of 48 described species represented) and low data matrix occupancy (4-46%, Table S2) of Cubozoa in our analyses (Fig. S2–S4) and Kayal et al. (2018).

### Staurozoa

The phylogenetic placement of staurozoans has been recalcitrant (Fig. S1). Historically, these so-called stalked jellyfish were considered scyphozoans until Marques and Collins (2004) elevated the group to the class level (Miranda, et al. 2010). Despite the variable position of Staurozoa in our phylogenies (see above), the relationships within the clade were constant and matched those of Kayal et al. (2018), the source of the transcriptomic data analyzed here. The relationships among the genera sampled here were discordant with those inferred by Miranda et al. (2016), which to date has the best taxon sampling for Staurozoa, based on a concatenated matrix of mitochondrial (COI, 16S) and nuclear genes (ITS, 18S, 28S) under parsimony, maximum likelihood, and Bayesian analyses.

Within Staurozoa, transcriptome-based analysis suggests Myostaurida gave rise to Amyostaurida. This result would suggest that muscles in the stalks have been lost in Amyostaurida (also suggested in Miranda, Collins, et al. 2016; Miranda, Hirano, et al. 2016; Miranda, et al. 2018) because scyphozoan polyps possess these muscles. However, the constrained 18S tree paints a more complicated picture, as most members of Amyostaurida diverge near the base of the clade and representatives of Myostaurida are split into two separate clades (Fig. 3).

### Scyphozoa

Relationships within Scyphozoa are consistent and highly supported across all our phylogenies (Fig. 2, S2–4) and agree with prior studies that found Discomedusae sister to Coronatae ((Bayha, et al. 2010; Bayha, et al. 2017). Despite removing *Periphylla periphylla* due to high numbers of transcript copies and/or paralogs (Table S2), we recovered the same relationships among the remaining 6 scyphozoan species as Kayal et al. (2018), the source of the transcriptomic data analyzed here. Interestingly, despite frequent topological discordance between nuclear and mitochondrial phylogenies, we find the same phylogenetic relationships (a clade containing *Cyanea* and *Chrysaora* sister to a clade containing *Aurelia* and *Cassiopea*) as Kayal et al. (2013), who inferred phylogenies using mitogenomes, and Daglio and Dawson (2017) who used mitochondrial and nuclear ribosomal genes (16S, 18S, 28S). Comparisons to other recent phylogenomic studies are limited due to low taxon sampling in this group (Zapata et al. 2015, 2 scyphozoans; Quattrini et al. 2020, 1 scyphozoan).

### Hydrozoa

The transcriptomic phylogeny (Fig. 2) recovered Hydrozoa split into Trachylina and Hydroidolina, in agreement with analyses of other previous studies (Collins, et al. 2008; Zapata, et al. 2015; Kayal, et al. 2018; Picciani, et al. 2018; Bentlage and Collins 2021), although the 18S topology recovered the trachyline *Halammohydra* as falling outside of these two clades, rendering Trachylina paraphyletic. The 18S topology fails to recover the monophyly of and relationships among many well-established taxa. For example, the 18S topology includes part of a grade of Limnomedusae (including Geryonidae) plus Actinulida as sister to the remainder of Hydrozoa, rendering Trachylina paraphyletic while Narcomedusae was monophyletic, derived from within the grade of Trachymedusae, which is congruent with previous studies (Collins, et al. 2008; Bentlage, et al. 2018) (including Actinulida). Within Hydroidolina, two distinctive clades (Siphonophorae and Leptothecata) were monophyletic, but Aplanulata, Capitata and Filifera were not. The latter was as expected given that earlier studies established Filifera as paraphyletic (e.g., Cartwright, et al. 2008; Bentlage and Collins 2021). The high degree of conservation of 18S and resulting lack of resolution from 18S has hampered inferences of hydrozoan relationships previously, an issue that has been partially overcome by using additional markers such as 16S and 28S (Collins, Bentlage, et al. 2006; Cartwright, et al. 2008; Collins, et al. 2008) and most recently target capture data (Bentlage and Collins 2021).

While the 18S topology failed to recover monophyly of most of the traditionally recognized groups within Hydrozoa (Fig. 3), the transcriptomic phylogeny (Fig. 2) was consistent with many previous studies. Within Hydroidolina, Aplanulata (*Ectopleura* and *Hydra*) is shown to be the sister to the rest of Hydroidolina, consistent with previous phylogenomic studies (Bentlage and Colllins 2021; Zapata et al., 2015, Kayal et al., 2018). In addition, Filifera III and IV (*Hydractinia, Podocoryne, Turritopsis*) were recovered as monophyletic (Fig. 2), consistent with Bentlage and Collins (2021), Cartwright et al. (2008), Kayal et al., (2018) and Zapata et al., (2015). Sampling of genomic data has been particularly sparse for Leptothecata, the most species rich of all roughly ordinal-level medusozoan clades (just one of 2,140 leptothecate species sampled herein) and Filifera I and II, which are not represented in the genomic data analyzed herein. Bentlage and Collins (2021) filled some of these gaps with target capture data but more sampling will be necessary to settle some of the long-standing debates in hydrozoan systematics. The sparse taxonomic representation of hydrozoans, and resulting small backbone may at least partially explain the lack of resolution of the 18S phylogeny for Hydrozoa.

### Future of phylogenomics for cnidarians and all other organisms

One critical need for future studies is to improve taxon sampling across the cnidarian tree. Currently, we have just scratched the surface of capturing the species richness of Cnidaria in our phylogenomic analyses (Fig. 1A). Of the 12,000 plus cnidarian species known, the largest phylogenomic study to date (Quattrini et al. 2020) includes just 2% of cnidarian species richness and our 18S hybrid phylogeny includes less than 15% (Fig. 1A). Recent studies have dramatically improved the representation of certain clades, for example Anthozoa and Octocorallia (Quattrini et al. 2017, 2020), and Actinaria (this study), while other clades like Medusoza have lagged (Fig. 1B), particularly from the species-rich hydrozoan clade Leptothecata. Cubozoans also have been poorly represented (Fig. 1b, Chang et al. 2015 *Tripedalia cystophora* only, Zapata et al. 2015 *Alatina alata* only, Stampar et al. 2019 *Alatina moseri* only, Quattrini et al. 2020 *Alatina alata* only), making relationships within the clade difficult to resolve with confidence. Zoantharia, Endocnidozoa, Antipatharia, and Ceriantharia are other groups that have historically been underrepresented in phylogenomic studies (Fig. 1B). Furthermore, sampling bias due to uneven geographic accessibility presents a major challenge in building a “complete” tree of Cnidaria.

The 18S phylogeny we estimated employed a hybrid approach leveraging the strength of the multi-locus phylogenomic backbone and the vast number of 18S sequences publicly available. This method is useful for identifying areas where taxonomy can be improved, although instances of contamination and misidentification in large databases like GenBank can produce spurious results in phylogenetic trees. In some cases, the topology based on 18S was unconventional, differing from results seen in focused analysis of this marker. This may reflect differences in alignment and models in a phylum-scale data set and highlight areas of the tree where molecular evolution differs from that of the clade as a whole. Our backbone phylogeny, estimated from orthologs mined from transcriptomes and gene models, corroborates relationships recovered in recent UCE phylogenies.

Target capture approaches like UCEs are likely the way forward in phylogenomic studies in cnidarians and other taxa, given their cost effectiveness, coverage, and flexibility, particularly in that they can be used on older and/or less optimally preserved specimens (McCormack, et al. 2016; Ruane and Austin 2017; Derkarabetian, et al. 2019). Target capture approaches also reduce problems with multiple transcript isoforms and paralogous loci, which we encountered mining transcriptomes for phylogenetic markers. Generating genome and transcriptome sequences will remain important however, since these data aid in the design of target capture probe sets (e.g., Quattrini, et al. 2018) and are required for other analyses (e.g., selection, structural variants, gene family evolution, among others).

### Conclusions

The goal of generating a cnidarian tree that includes nearly every species increasingly seems like a reality that will happen within the next 10 or 20 years. Nevertheless, generating trees (i.e., hypotheses of relationships) that encompass as many species as possible is critical for moving the field of cnidarian systematics and supplying cnidarian researchers with a framework from which to interpret evolutionary patterns. Our transcriptome-based phylogenies confirm backbone relationships resolved in recent studies using target capture, which is a promising approach for reaching the goal of total taxon sampling. Until we reach that goal, our 18S hybrid phylogeny increases taxon sampling to almost 15% of the total cnidarian species richness, providing a roadmap for taxonomy and trait evolution.

### Additional

## Acknowledgements

We would like to acknowledge the many researchers who generated and/or analyzed the public genomic, transcriptomic, and 18S data used in these analyses. We thank Gustav Paulay and Jessie Whelpley for sample collection and RNA extraction for *Leptogorgia* sp. and Renato Nagata for sample collection of *Arachnanthus* sp. We thank Antonis Rokas for discussions about phylogenomic analyses and Barbara Battelle for her 30-year leadership of the Whitney Lab REU program. We would like to acknowledge David Plachetzki for input on this manuscript. The color palette of figures was inspired by *The Warmth of Other Sons* by Bisa Butler, 2020.

## Supporting Information

All analysis scripts, alignments, and trees are available at https://github.com/josephryan/DeBiasse_cnidophylogenomics and transcriptomes are available on Dryad (https://doi.org/10.6071/M3K39S). A snapshot of the GitHub repo is included here as supplemental file 1. All accessions are available in Table S2.

## Supplementary information

**Figure S1.**
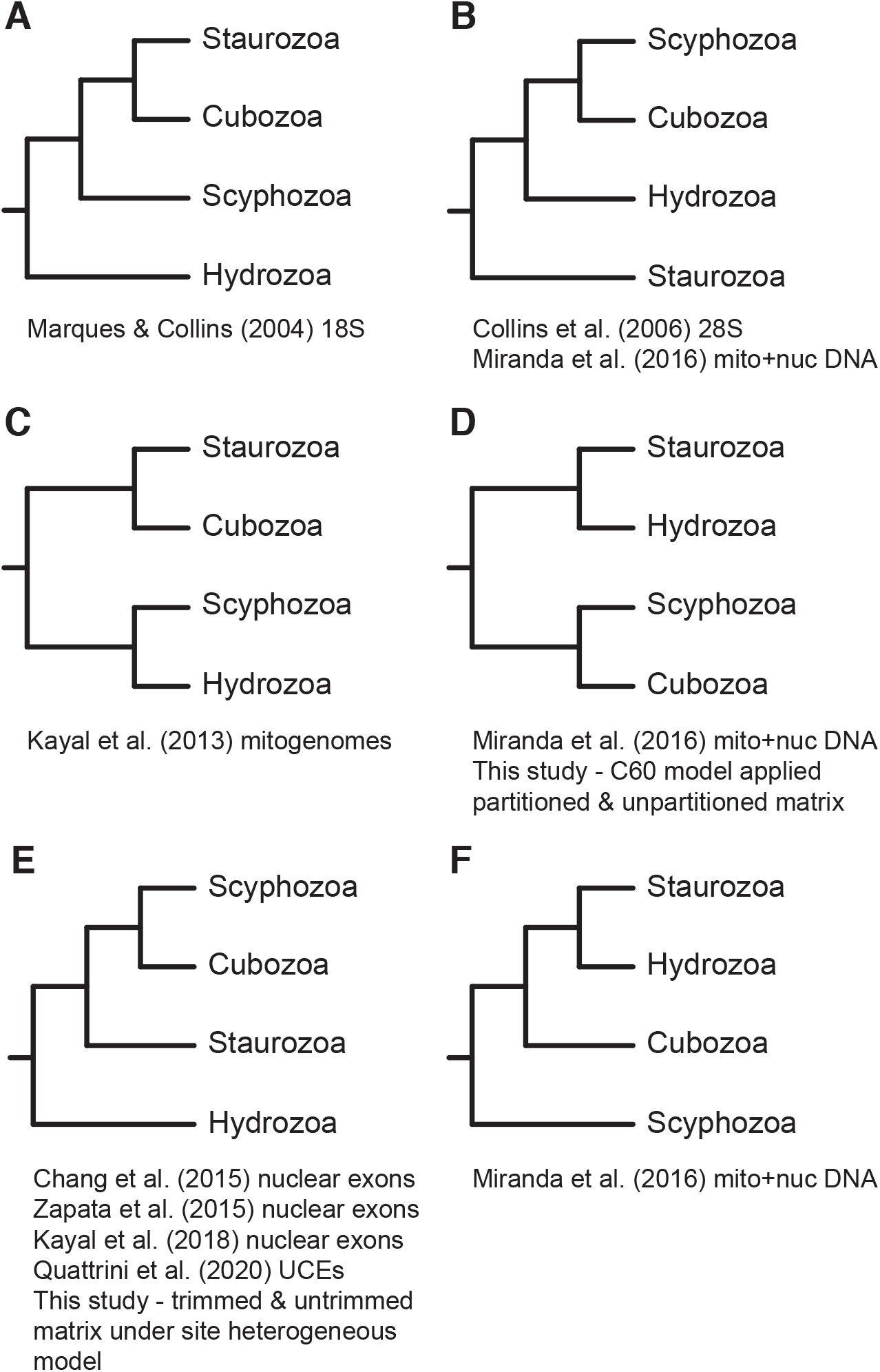
Cladograms representing the historical phylogenetic relationships found within Medusozoa

**Figure S2.**
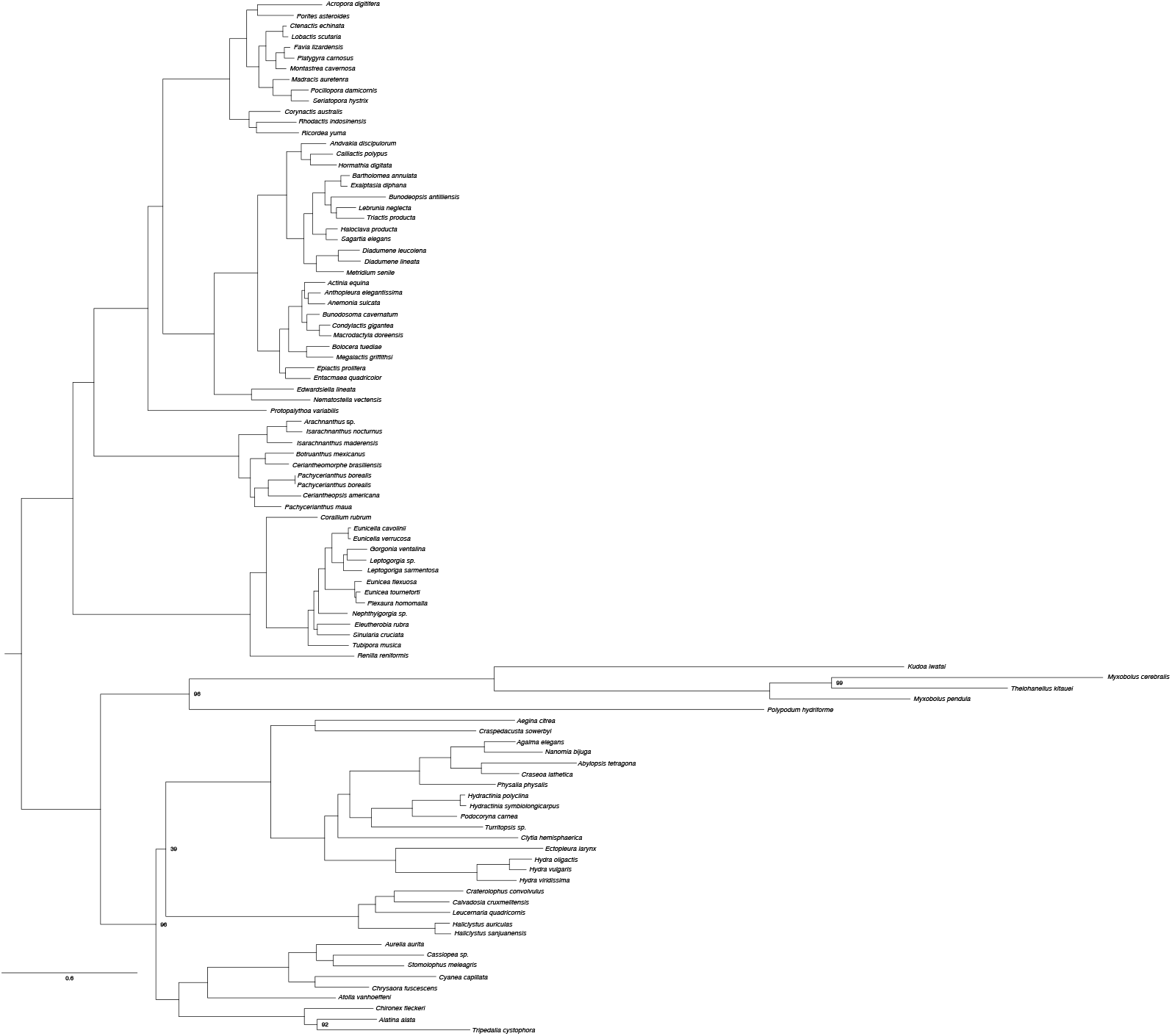
Maximum-likelihood phylogeny of cnidarian estimated from a concatenated matrix of 748 trimmed ortholog alignments generated partition-specific models. Bootstrap values are indicated at nodes with less than 100% support.

**Figure S3.**
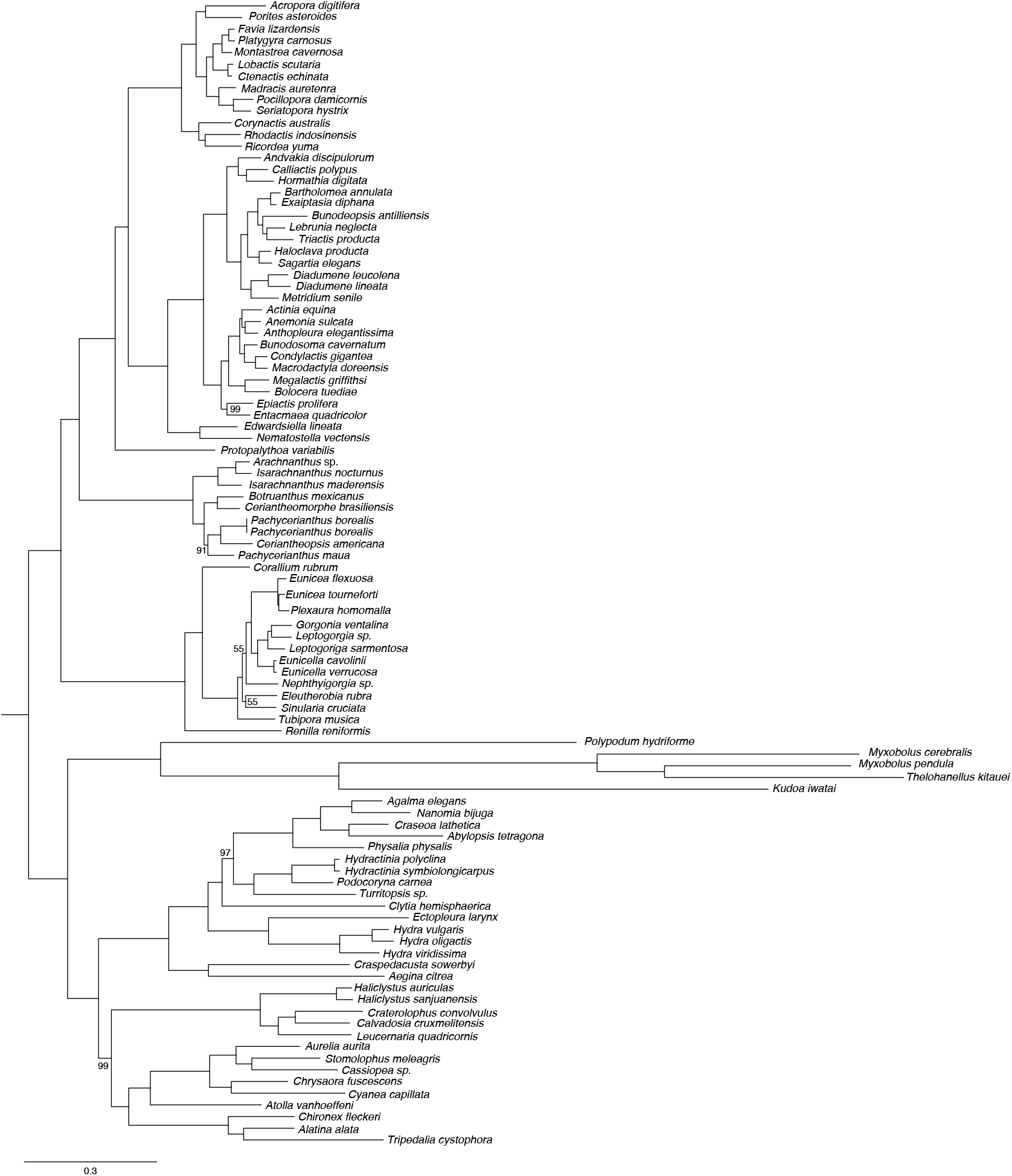
Maximum-likelihood phylogeny of cnidarians estimated from a concatenated matrix of 748 untrimmed ortholog alignments under the C60 model of amino acid substitution. Bootstrap values are indicated at nodes with less than 100% support.

**Figure S4.**
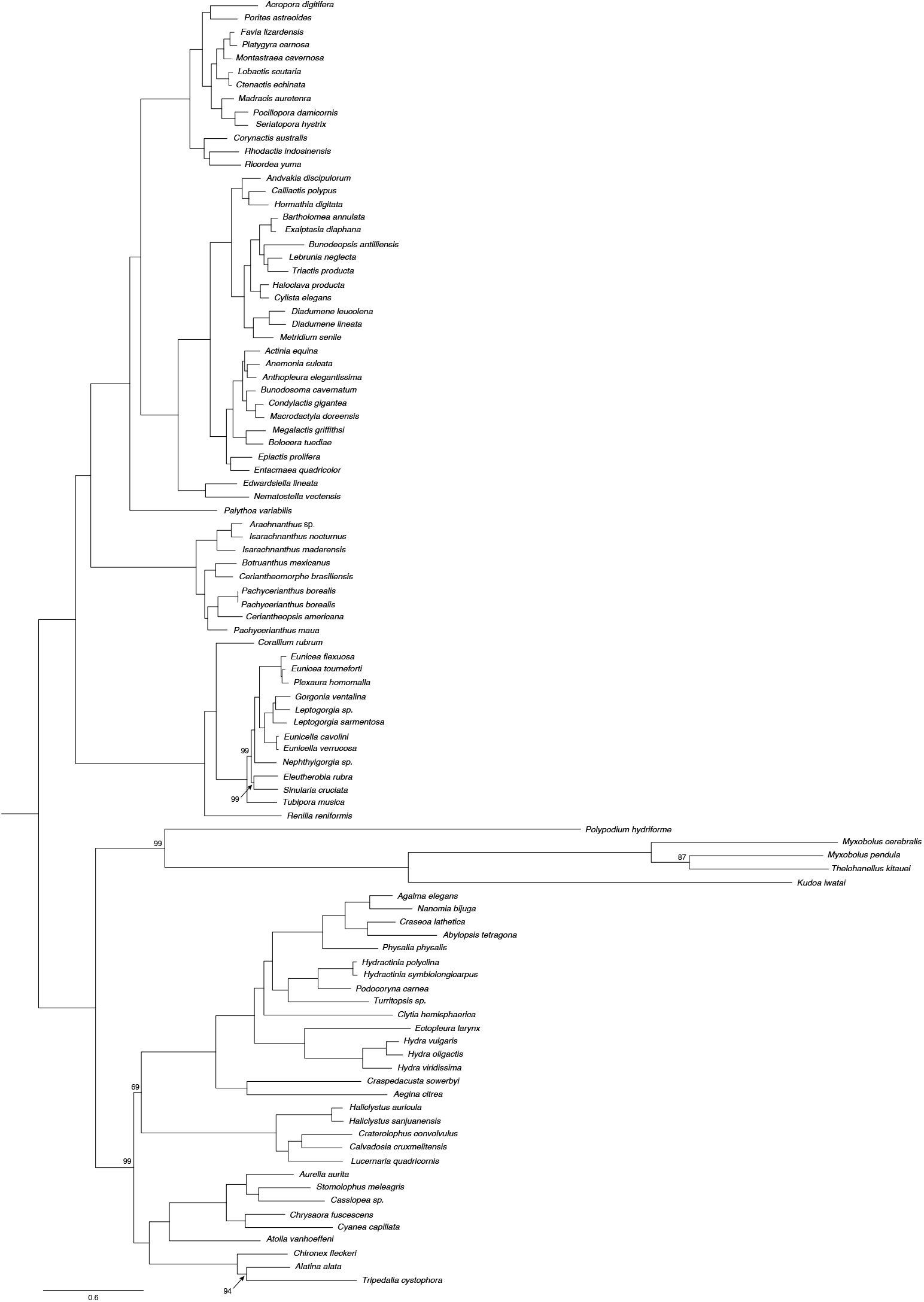
Maximum-likelihood phylogeny of cnidarians estimated from a concatenated matrix of 748 untrimmed ortholog alignments under the C60 model of amino acid substitution applied to a partitioned matrix. Bootstrap values are indicated at nodes with less than 100% support.

